# ROS-Induced Stress Promotes Enrichment and Emergence of Antibiotic Resistance in Conventional Activated Sludge Processes

**DOI:** 10.1101/2024.04.05.585878

**Authors:** Bharat Manna, Xueyang Zhou, Naresh Singhal

**Affiliations:** Department of Civil and Environmental Engineering, University of Auckland, Auckland 1142, New Zealand; Water Research Centre, University of Auckland, Auckland 1142, New Zealand

**Author notes:** Corresponding Author: Naresh Singhal.

**Keywords:** wastewater treatment, great oxygenation event, reactive oxygen species, metagenomics, SWATH-metaproteomics

## Abstract

Since the Great Oxidation Event 2.4 billion years ago, microorganisms have evolved sophisticated responses to oxidative stress. These ancient adaptations remain relevant in modern engineered systems, particularly in conventional activated sludge (CAS) processes, which serve as significant reservoirs of antibiotic resistance genes (ARGs). While ROS-induced stress responses are known to promote ARG enrichment/emergence in pure cultures, their impact on ARG dynamics in wastewater treatment processes remains unexplored. Shotgun-metagenomics analysis of two hospital wastewater treatment plants showed that only 35-53% of hospital effluent resistome was retained in final effluent. Despite this reduction, approximately 30% of ARGs in CAS showed higher abundance than upstream stages, of which 22% emerged *de novo*. Beta-lactamases and efflux pumps constituted nearly 50% of these enriched ARGs. These ARGs exhibited significant correlations (p < 0.05) with ROS stress response genes (*oxyR*, *soxR*, *sodAB*, *katG* and *ahpCF*). The CAS resistome determined 58-75% of the effluent ARG profiles, indicating treatment processes outweigh influent composition in shaping final resistome. Proof-of-concept batch reactor experiments confirmed increased ROS and ARG levels under high dissolved oxygen (8 mg/L) compared to low oxygen (2 mg/L). Untargeted metaproteomics revealed higher expression of resistant proteins (*e.g.*, OXA-184, OXA-576, PME-1, RpoB2, Tet(W/32/O)) under elevated ROS levels. Our findings demonstrate that CAS processes actively shape effluent resistome through ROS-mediated selection, indicating that treatment processes, rather than initial wastewater composition, determine final ARG profiles. This study indicates that the emergence of ARGs needs to be considered as an integral aspect of wastewater treatment design and operation to prevent antibiotic resistance dissemination.

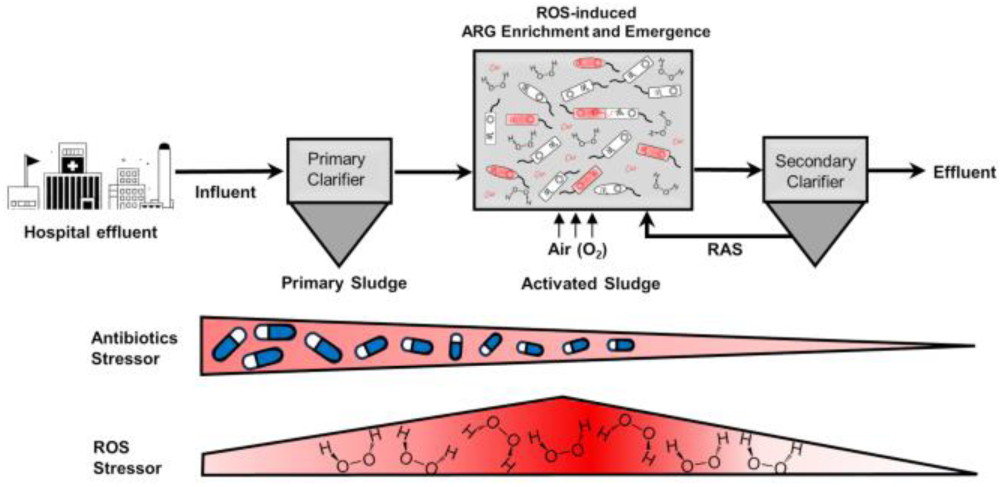

**Highlights:**

- Approximately 30% of ARGs enriched in CAS versus upstream stages, of which 22% emerged *de novo*
- Beta-lactamases and efflux pumps comprise ∼50% of enriched ARGs in CAS
- CAS resistome shapes 58-75% of effluent ARGs, outweighing influent composition
- High DO levels increase ROS production and ARG abundance in CAS systems
- Metaproteomics confirms higher expression of resistant proteins under elevated ROS levels

## 1. Introduction

Antibiotic resistance (AR) poses an escalating threat to global public health, responsible for approximately 1.27 million fatalities worldwide and contributing to almost an estimated 4·95 million (3·62–6·57) deaths associated with bacterial antimicrobial resistance in 2019 (Murray, Christopher JL, 2022). In the United States alone, infections afflict over 2.8 million individuals annually, leading to the deaths of more than 35,000 people (Centers et al., 2019). The financial burden of managing infections caused by six prevalent antibiotic-resistant bacteria (ARB) (Murray, Christopher JL, 2022) especially within healthcare settings exceeds $4.6 billion each year (Nelson et al., 2021). Antibiotic resistance is a critical One Health issue (McEwen and Collignon, 2018), profoundly affecting humans, animals, and the environment, impacting even the most fundamental resource for life: water.

The presence of antibiotic-resistant genes (ARGs) and ARBs often can be attributed to effluents from hospitals, farms, and sewage systems, illustrating the interconnectedness of human activities and the broader environmental impact of AR (Agga et al., 2015; Rowe et al., 2017). As wastewater treatment plants (WWTPs) operate at the human-environment interface, they play a crucial role in both managing and understanding the global health effects of AR (Alexander et al., 2020; Manaia et al., 2018). WWTPs receive and process diverse influents, including hospital effluents laden with high concentrations of antibiotics, ARBs, and ARGs (Y. Chen et al., 2020; Uluseker et al., 2021; Wang et al., 2022). This influx creates a potent selection pressure for ARGs and facilitates horizontal gene transfer (HGT) among bacterial communities (Karkman et al., 2018; Uluseker et al., 2021; Wei et al., 2021). Notably, hospital wastewaters harbor ARG concentrations several orders of magnitude higher than municipal wastewater (Korzeniewska et al., 2013), highlighting their alarming potential to seed and propagate resistance within WWTPs treating such influents (Hocquet et al., 2016; Mehanni et al., 2023; Paulus et al., 2019). Moreover, recent studies reported that different WWTP treatment processes could exhibit varying impacts on ARG and ARB dynamics (Honda et al., 2023; Luo et al., 2021; Majeed et al., 2021). Interestingly, the authors identified significant alteration in the ARGs composition in the sludge compared to that of the influent resistome and suggested seasonality in antimicrobial use to be key influencing factor behind this transition (Honda et al., 2023). A multi-omics study revealed significant enrichment of both absolute and relative abundance of specific resistance genes in CAS resistome compared to influent, particularly those conferring resistance to ARGs drug classes (multidrug, beta-lactam, aminoglycoside, tetracycline, vancomycin etc.), suggesting bacterial community composition and biomass concentration as key factors (Ju et al., 2019). Moreover, another study showed that while the overall ARGs abundance decreased during 11 out of 12 WWTPs, there were instances of specific ARGs being enriched in the effluents (Garner et al., 2024). Many other reports also identified CAS processes as potential hotspots for ARG enrichment (Bengtsson-Palme et al., 2016; Guo et al., 2017; Liu et al., 2018, 2019; Majeed et al., 2021; Petrovich et al., 2020). These enriched ARGs in CAS contribute significantly to the effluent resistome which is disseminated into the receiving environment (Honda et al., 2023; Ju et al., 2019; Majeed et al., 2021). However, the specific mechanisms driving the enrichment and emergence of ARGs within CAS processes remain poorly understood. Consequently, understanding the underlying mechanisms behind the emergence of ARGs in CAS processes has become an urgent need.

While antibiotics are relatively recent human inventions, microbes have developed resistance mechanisms over thousands of years through their continuous evolutionary struggle for survival (Dcosta et al., 2011). To understand bacterial resistance mechanisms, the perspective of oxygen on Earth provides a critical evolutionary context. Prior to the Great Oxidation Event (GOE) around 2.4-2.1 billion years ago, Earth’s atmosphere was essentially oxygen-free, and early life thrived in a reducing condition (Lyons et al., 2014). The dramatic rise of oxygen through photosynthesis represented the first major oxidative stress that organisms had to adapt to, driving the evolution of cellular mechanisms to cope with oxygen toxicity and its reactive species (ROS). Recent evidence suggests a unifying mechanism behind the action of many bactericidal antibiotics, where ROS generation plays a key role behind lethality (Dwyer et al., 2009; Lam et al., 2020; Van Acker and Coenye, 2017). ROS like hydrogen peroxide (H_2_O_2_), hydroxyl radical (OH^·^), singlet oxygen (^1^O_2_), and superoxide anion (O_2_^⁻^) are inherent byproducts of bacterial aerobic respiration (Imlay, 2019, 2003). Furthermore, ROS-mediated killing can trigger a secondary stress response beyond the initial antibiotic-induced damage, involving factors like activation of response regulators (*e.g.*, *ArcA*), metabolic reprogramming of the TCA cycle, and disruption of the nucleotide pool with increased ATP demand (Li et al., 2021) (Figure S1). To counter this oxidative stress and survive, bacteria have evolved diverse antioxidant defenses through metabolic remodeling (Li et al., 2021). However, bacterial efforts to develop ROS tolerance often come at a cost. The increased oxidative stress can lead to direct DNA damage, potentially triggering novel mutations, and ultimately contributing to the emergence of AR (Ariza et al., 1994; Jin et al., 2018; Mandsberg et al., 2009; Qi et al., 2023). Since ROS are common to both antibiotic action and bacterial stress responses, resistance to ROS could be a key shared characteristic underlying ARG development. We recently showed that ROS stress has been linked to ARGs proliferation in freshwater microbial communities (Manna et al., 2024).

These ancient adaptations to oxidative stress remain fundamental to how modern microorganisms handle ROS in highly aerated wastewater treatment systems. The significance of ROS in CAS processes can be attributed to the constant aeration inherent to these systems, which creates a unique environment characterized by high dissolved oxygen (DO) levels and, consequently, elevated ROS generation. While existing research hints at the connection between ROS and ARG emergence in the context of WWTP, it primarily focuses on lab experiments and specific antibiotics or stressors. One study observed a significant increase in antibiotic resistance in *Escherichia coli* exposed to aeration tanks compared to primary or final sedimentation tanks (Sulfikar et al., 2018). Another study linked triclosan-induced oxidative stress to a rise in ARGs, particularly tetracycline and multidrug resistance genes (Tan et al., 2021). Similarly, ROS generated by phenolic compounds have been proven to promote resistance through increased membrane permeability and HGT (Ma et al., 2021). Moreover, antidepressant fluoxetine induced resistance to multiple antibiotics (*e.g.*, fluoroquinolone, aminoglycoside, β-lactams, tetracycline and chloramphenicol) in *Escherichia coli* via ROS-mediated mutagenesis (Jin et al., 2018). Despite these findings, the critical question of whether aeration-induced ROS in real-world WWTPs shape the multifaceted resistome in CAS processes, remains unanswered.

Building on this evolutionary and mechanistic foundation, we present a comprehensive investigation of how aeration-induced oxidative stress shapes antibiotic resistance in full-scale CAS processes. By combining whole-genome shotgun metagenomics analysis from two WWTPs treating hospital effluents and untargeted metaproteomics from controlled batch reactor experiments, we unravel the relationship between DO levels, ROS generation, and ARG dynamics, providing unprecedented insights into how oxidative stress drives resistance development in CAS processes.

## 2. Materials and methods

### 2.1. Sample data collection

In this work, we considered paired-end shotgun metagenomics samples obtained using a HiSeq 2500 instrument from two full-scale WWTPs processing university hospital effluents in Göttingen (WWTP_Gö_), and Greifswald (WWTP_Gw_), Germany (Schneider et al., 2020). The samples were collected from five different stages (abbreviated as S) from both the WWTPs, *i.e.*, S1: Hospital effluent (a storage tank next to the university hospital), S2: WWTP Influent, S3: Primary Sludge, S4: conventional activated sludge (CAS), and S5: WWTP effluent as described elsewhere (Schneider et al., 2021, 2020). The details of the sample names, and NCBI SRA accession used in this study, can be found in Table S1 and Table S2. The raw metagenomes can be accessed in the NCBI Sequence Read Archive under the BioProject accession number PRJNA524094 (Schneider et al., 2021).

### 2.2. Metagenomics Analysis

We performed preprocessing steps to eliminate low-quality sequences using Trimmomatic v0.39 (Bolger et al., 2014) with specific quality thresholds (Manna et al., 2024). For metagenomic analysis, we employed the SqueezeMeta v1.5.2 pipeline (Tamames and Puente-Sánchez, 2019). The read statistics per sample are provided in Table S1 and Table S2. Co-assembly was performed using Megahit v1.2.9 (D. Li et al., 2015). Within the SqueezeMeta pipeline, Barrnap v0.9 (Seemann, 2014) tool was used for predicting RNAs, while Prodigal v.2.6.3 (Hyatt et al., 2010) was utilized for predicting ORFs. Moreover, the Comprehensive Antibiotic Resistance Database (Alcock et al., 2020) release 3.2.4 was used for annotating the ARGs using Diamond v.2.0.14 (Buchfink et al., 2014) with sequence identity of 70% and e-value threshold of 1 × 10^−3^ (Manna et al., 2024; Petrovich et al., 2020). ARG abundances were normalized using the 16S-rRNA gene abundance as per previous report (B. Li et al., 2015). ARGs conferring resistance to more than one antibiotic was classified as ‘multidrug resistant’ (MDR). Similarity searches against Kyoto Encyclopedia of Genes and Genomes (Kanehisa and Goto, 2000), were performed for functional assignments of ORFs. The metagenomics data was analyzed using the SQMtools v1.6.3 R package (Puente-Sánchez et al., 2020).

### 2.3. Bioinformatics Analysis and Visualization

To assess the diversity of ARG composition across the different treatment stages, we calculated a Bray-Curtis dissimilarity matrix with PERMANOVA to determine the significance of clustering, using MicrobiomeAnalyst v2.0 (Dhariwal et al., 2017). A heatmap was plotted to visualize changes in the abundance of each ARG, utilizing a z-score with TBtools v2.007 (C. Chen et al., 2020). Furthermore, differentially abundant (enriched/emerged) ARGs in CAS (S4) were identified using EdgeR v.4.4.0 (Robinson Mark et al., 2010) with an adjusted *p*-value cutoff of 0.05. The correlation between ROS stressor genes and differentially abundant ARGs was performed using a Mantel test (*p*-value < 0.05) using OmicStudio tools v1.41.0 (Lyu et al., 2023). Graphics were generated using the ggplot2 v3.5.1 (Wickham, 2011) package in R (R Core Team, 2022) version 4.2.1.

### 2.4. Batch reactor experiments with different DO levels

To validate our metagenomics-based findings of higher ROS generation under more oxidative stress leading to increased ARGs, we performed batch reactor experiments with CAS samples under low (2 mg/L) and high (8 mg/L) DO levels. The CAS samples were obtained from the Māngere WWTP in Auckland, New Zealand. The experiments were conducted in reactors (n = 3 biological replicates per condition) with a capacity of 1 L, at a temperature of 20 °C for 48 hours. Composition of the synthetic wastewater is provided in Table S3 and Table S4. Detailed methodology for ROS measurement, DNA extraction for metagenomics analysis and untargeted metaproteomics analysis are provided in Supplementary Methods. The raw metagenome samples from these experiments can be accessed in the European Nucleotide Archive under the BioProject accession number PRJEB74089 as provided in Table S5.

### 2.5. SWATH-MS based quantitative Metaproteomics Analysis

Untargeted metaproteomic analysis was performed on activated sludge samples to characterize microbial protein expression under the two DO conditions. Proteins were extracted using SDS-based lysis buffer, followed by sonication and TCA-acetone precipitation. Tryptic-digested peptides were analyzed using a NanoLC 400 UPLC (Eksigent, USA) system coupled to a TripleTOF 6600 mass spectrometer (Sciex, USA). Protein identification using MetaProteomeAnalyzer v3.4 (Muth et al., 2015) with X-tandem revealed 1,110 distinct proteins for functional analysis. Detailed methodology is available in Supplementary Methods. The metaproteomics data has been submitted to ProteomeXchange Consortium with the dataset identifier PXD044490.

## 3. Results and Discussion

### 3.1. Comparison of ARG abundances across different treatment stages

In this study, we systematically analyzed and compared metagenomic samples from various stages of wastewater treatment processes to assess changes in the resistome profile. These stages included hospital effluents (S1), WWTP influent (S2), primary treatment (S3), CAS (S4), and WWTP effluents (S5) (Figure 1a). Our findings showed a considerable decrease in total ARG abundances at different stages of WWTP_Gö_ compared to the hospital effluents: 54.17% in S2, 53.38% in S3, 78.69% in S4, and 83.30% in S5 (Figure 1b). Moreover, 69.50%, 68.42%, 46.59%, and 35.14% of the ARGs detected in S1 were retained in S2, S3, S4 and S5, respectively (Figure S2a).

**Figure 1.**
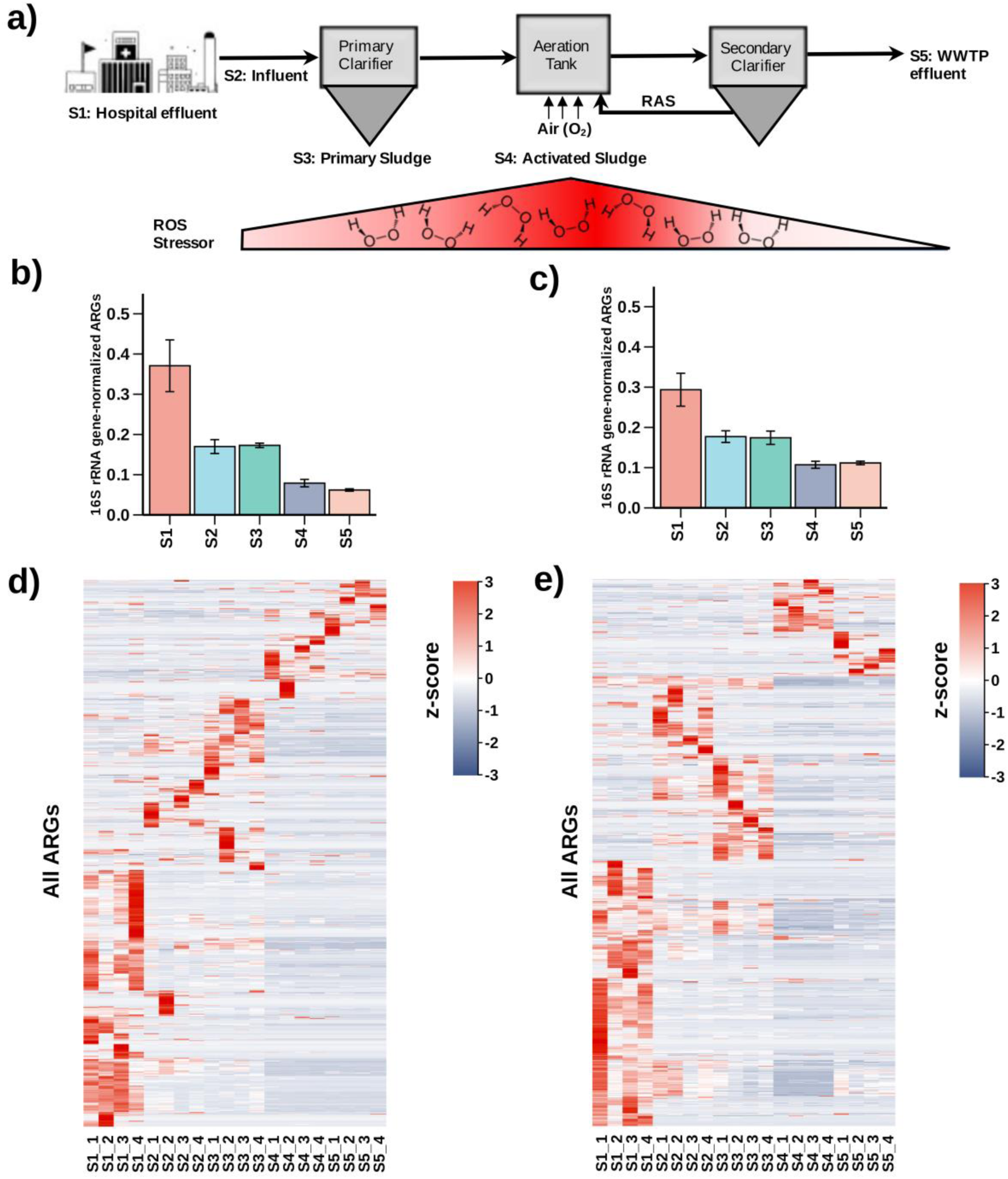
Alteration of ARG abundance across different WWTP stages. (a) Schematic representation of the WWTP stages studied, including S1: Hospital effluent, S2: WWTP influent, S3: Primary sludge, S4: Activated sludge, and S5: WWTP effluent. The variation in ROS stressor levels across these stages is depicted below. (b) and (c) show the total relative ARG abundance at each stage for WWTP_Gö_ and WWTP_Gw_, respectively. Heatmaps display the trend of change in each ARG relative abundance across the stages in (d) WWTP_Gö_ and (e) WWTP_Gw_ with a prominent set of enriched (or emerged) ARGs (red) in both S4 and S5. RAS: Recycled activated sludge.

A similar analysis conducted in the WWTP_Gw_ also revealed a reduction in ARG abundances of 39.70% in S2, 40.65% in S3, 63.55% in S4, and 61.99% in S5, compared to S1 (Figure 1c). The numbers of ARGs detected in S1 were retained by 73.14% in S2, 67.55% in S3, 31.52% in S4, and 53.42% in S5 (Figure S2c). The highest abundance of ARGs was observed in the hospital effluent (S1) for both WWTPs, which is expected due to the prevalent use of antibiotics in hospital settings, leading to a higher presence of ARGs. Across the treatment stages, a consistent trend of ARG reduction was noted, indicating the effectiveness of the WWTP processes in decreasing the overall ARG abundance in the effluent (S5). This pattern of ARG reduction in CAS and effluents are in agreement with previous metagenomics-based studies (Bengtsson-Palme et al., 2016; Honda et al., 2023; Lira et al., 2020).

Furthermore, the diversity of resistome profiles across various stages of the WWTP_Gö_ and WWTP_Gw_ was evaluated using principal coordinate analysis (PCoA) with a Bray-Curtis dissimilarity matrix (Figure S2b, d). The PCoA demonstrated clear clustering of resistomes corresponding to the five treatment stages. Significant separation observed (R^2^: 0.79329, PERMANOVA F-value: 14.392, *p*-value: 0.001) in WWTP_Gö_, indicating distinct ARG profiles at each treatment stage. Similarly, WWTP_Gw_’s resistome profiles were markedly segregated demonstrating substantial differentiation (R^2^: 0.83563, PERMANOVA F-value: 19.064, *p*-value: 0.001). The PCoA results showed that the resistomes from S1, S2, and S3 stages clustered closely, suggesting similar ARG profiles, possibly due to the direct contribution of hospital effluent to the influent. In contrast, samples from the CAS (S4) and treated effluent (S5) stages were distinctly separate from the preceding stages. This delineation reflects a significant transition in the ARG profile, attributable to the biological processes and selection pressure within the activated sludge system and secondary clarifiers. Notably, the effluent resistome consisted of 58.42% and 75.58% of the CAS resistome in the WWTP_Gö_ and WWTP_Gw_, respectively, indicating a substantial overlap between the systems. The distinct clusters for the S4 and S5 stages also suggest that the treatment process, rather than initial wastewater composition, determines final effluent ARG profiles, with potential environmental implications upon its release. Similar variations in resistome in influent, activated sludge and effluent have been reported in previous studies (Honda et al., 2023; Majeed et al., 2021).

In addition to the overall trend analysis, a detailed heatmap of all ARGs based on normalized abundance (z-scores) was generated to further elucidate the specific changes in ARG profiles at each treatment stage, with a focus on the CAS (S4) process (Figure 1d, e). The heatmap illustrated that while some ARGs were consistently abundant across all stages, others were more stage specific. This abundance pattern reinforces the notion of stage-specific selective pressures and microbial community dynamics shaping the ARG profile. Interestingly, despite the general trend of decreasing total ARG abundance (Figure 1b, c), certain ARGs exhibited notable persistence and remarkable enrichment/emergence in the CAS stage (S4) in both the WWTPs (Figure 1d, e). This relative enrichment of ARGs in the CAS was previously reported using combined metagenomics and metatranscriptomics study (Ju et al., 2019).

### 3.2. Identifying the enriched and emerged ARGs in the CAS process

To enhance our understanding of the enriched or emerged ARGs within the CAS process, we conducted a detailed analysis of ARGs that were differentially more abundant in CAS than in the upstream stages. Specifically, in the WWTP_Gw_ facility, we observed that 111, 115, and 116 ARGs exhibited significantly (FDR < 0.05) higher abundance in CAS when compared to S1, S2, and S3, respectively, as illustrated in Figure 2a. Similarly, for the WWTP_Gö_ facility, 152, 138, and 114 ARGs were found to be more abundant in CAS relative to the same comparator stages (Figure S3a). Additionally, certain ARGs were identified as being significantly more abundant in S4 compared to all other stages (S1, S2, and S3), with some ARGs being uniquely abundant to specific treatment stages (Figure S4 and Supplemental Data 1). To facilitate downstream analyses, we compiled a list of ARGs that were significantly abundant in the CAS process, selecting those genes that displayed a higher average abundance in CAS compared to all preceding treatment stages. This analysis yielded a total of 116 (29.59%) and 94 (36.43%) highly abundant ARGs for WWTP_Gö_ and WWTP_Gw_, respectively, as detailed in Supplemental Data 2a. Notably, 22.22% (WWTP_Gö_) and 21.88% (WWTP_Gw_) of the abundant ARGs were not detected at any of the upstream stages, suggesting they emerged under the selection pressure of the CAS process (Supplemental Data 2b). These emerged ARGs exhibited resistance to aminoglycoside (*AAC(3)-IIIc*, *AAC(6’)-Iih*), beta-lactam (*OXA-34*, *OXA-660*, *OXA-4*, *GOB-3*, *GOB-4*, *IDC-2*, *IMP-1*, *IMP-13*, *IMP-28*), rifamycin (*iri*, *arr-1*), tetracycline (*tap*, *tet(V)*), glycopeptide (*vanD*, *vanI*), and other antibiotics. Additionally, among the enriched ARGs in both WWTPs were those belonging to various beta-lactamases, such as *OXA* (including *OXA-45*, *OXA-101*, *OXA-129*, *OXA-198*) beta-lactamases. These ARGs confer resistance to antibiotics like carbapenems, cephalosporins, and penams, highlighting a significant selection pressure for beta-lactam resistance. This is especially concerning given the extensive use of beta-lactam antibiotics and the potential for these resistance mechanisms to spread into the environment. Furthermore, we identified multiple ARGs conferring resistance to other antibiotic classes, including tetracycline (*tet(35)*, *tet(49)*, *tet(G)*, *tet(T)* etc.), glycopeptide (*vanC*, *vanF*, *vanS_in_vanC_cl*, etc.), and aminoglycoside (*AAC(3)-Id*, *AAC(6’)-32*, *AAC(6’)-Ic*, *AAC(6’)-IIa*, *AAC(6’)-IIb*, *AAC(6’)-IIc*, etc.), among others. Resistance to rifamycin (*rpoB2*, *Bado_rpoB_RIF*, *Nfar_rox*, *rphA*, *rphB,*), macrolide (*EreB*, *mphA*), fluoroquinolone (*QepA4*, *QnrS6*, *QnrS8*), and sulfonamide (*sul2*, *sul4*) antibiotics were also observed, alongside ARGs resistant to multiple antibiotics, such as *AxyY*, *efrB*, *MexF*, *mexP*, *mexQ*, *mexY*, *opmE*, *smeE*, and more, as provided in Supplemental Data 2a. Amongst the enriched/emerged ARGs, beta-lactamases (WWTPGö: 24.14%, WWTPGw: 25.53%) and efflux pumps (WWTPGö: 23.28%, WWTPGw: 27.66%) represented the predominant resistance mechanisms in both WWTPs. Our recent study also demonstrated that oxidative stress conditions could exacerbate ARG proliferation, particularly efflux pump genes (Manna et al., 2024). Consistent with our observations, a significant enrichment of various resistance genes, including those encoding for beta-lactamases, macrolides, and tetracyclines, was identified in recirculated activated sludge in comparison to primary sludge (Bengtsson-Palme et al., 2016). This enrichment was suggested due to the selective pressure within the wastewater treatment facility. Similarly, previous research documented an escalation in the relative abundance of a wide array of ARGs, spanning multiple drug classes such as beta-lactamases, aminocoumarins, glycopeptides, rifamycins, and sulfonamides, from influent to effluent in a conventional WWTP (Majeed et al., 2021). Furthermore, another study reported an increase in both the transcript and gene abundance of various ARG classes, including Multidrug, beta-lactamases, aminoglycosides, tetracyclines, trimethoprim, and vancomycin, in secondary effluent compared to the influent of post-primary clarifiers (Ju et al., 2019). Additionally, it was found that aerobic biomass exhibited a higher overall abundance of ARGs compared to anaerobic and anoxic biomass, with a notable increase in effluent abundances of fluoroquinolone and tetracycline ARGs (Petrovich et al., 2020). These observations underscore the diverse spectrum of resistant ARGs emerging within the CAS process, indicating that while the CAS process effectively reduces the overall ARG load, it also inadvertently facilitates the proliferation (enrichment and emergence) of multiple concerning ARGs.

**Figure 2.**
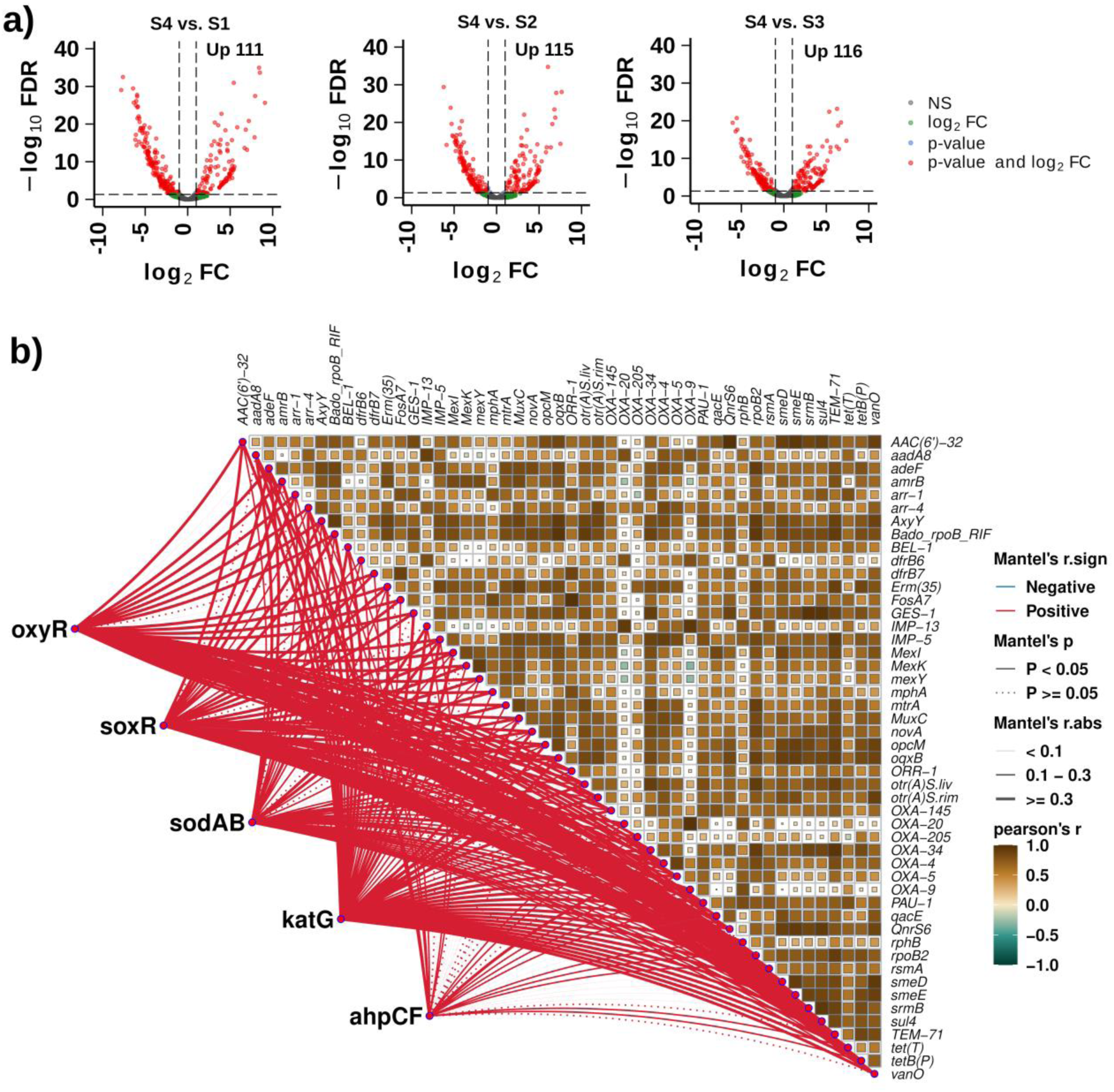
Differentially enriched (or emerged) ARGs and their correlation network with ROS stressors in the CAS process. a) Volcano plots illustrate the differentially abundant ARGs in the CAS stage compared to S1, S2, and S3 in the WWTP_Gw_ (log_2_FC ≥ 1, FDR < 0.05). b) A network analysis depicting the correlations between selected enriched/emerged ARGs and ROS stress response genes in CAS process in WWTP_Gw_.

### 3.3. Assessing the correlation between the ROS stressors and enriched or emerged ARGs

To further assess previously established connections between oxidative stress and antibiotic resistance, we analyzed correlations between ROS-response genes and the enriched/emerged ARGs detected in the CAS systems of both WWTPs. We focused on five crucial ROS-related genes (Seo et al., 2015): two transcription factors, *oxyR* and *soxR*, which play key roles in the cellular response to oxidative stress, and three genes encoding enzymes that help detoxify ROS, namely *sodAB* (Superoxide dismutase), *katG* (Catalase-peroxidase), and *ahpCF* (alkyl hydroperoxide reductase). Notably, previous reports revealed *oxyR* (Lau et al., 2005; Wei et al., 2012) and *soxRS* (P. Wang et al., 2020) contributed to virulence in *E. coli*, *Pseudomonas aeruginosa*, and *Salmonella enterica*. Our analysis revealed a significant correlation between these ROS stress response genes and numerous ARGs in both WWTPs (Figure 2b, Figure S3b, Supplemental Data 3). Specifically, in the WWTP_Gö_, the number of ARGs positively correlated with each ROS stressor were *oxyR* (7), *soxR* (35), *sodAB* (22), *katG* (31), and *ahpCF* (17). Meanwhile, in WWTP_Gw_, the numbers of ARGs linked to the same stressors were *oxyR* (60), *soxR* (49), *sodAB* (52), *katG* (79), and *ahpCF* (17). In the case of WWTP_Gö_, we identified several ARGs showing correlations with different combinations of ROS stressors (Figure S5a). Most notably, 4 ARGs (*AAC(6’)-IIb*, *lmrC*, *OXA-4*, and *QepA4*) correlated with all ROS stressors examined. The largest individual correlations were observed with *soxR*, which correlated with 10 ARGs, including tetracycline resistance genes (*tet(T)*, *tet(W/32/O)*), and various others (*Abau_AbaQ*, *dfrG*, *HelR*, *mexN*, *otr(A)S.liv*, *TriC*, *vgaALC*). Furthermore, a significant group of 6 ARGs, including metallo-beta-lactamases (*IMP-13*, *IMP-28*), beta-lactamases (*OXA-9*), and others (*aadA8*, *arr-1*, *vanS_in_vanO_cl*), showed correlation specifically with both *sodAB* and *katG*, indicating a potential link between these antioxidant enzymes and antibiotic resistance mechanisms. Additional notable correlations were observed with *katG* alone, which correlated with 9 ARGs, including various resistance mechanisms (*adeF*, *dfrB2*, *mphA*, *MYO-1*, *OXA-198*, *smeE*, *vanD*, *vanS_in_vanP_cl*, *vgaD*), suggesting a broad spectrum of antibiotic resistance mechanisms being influenced by oxidative stress responses. Besides, within the WWTP_Gw_, 10 ARGs were correlated with all ROS stressors, including genes responsible for multidrug resistance (*adeF*), beta-lactamase activity (*OXA-4*, *OXA-5*, *PAU-1*, *IMP-5*), tetracycline resistance (*tet(T)*, *tetB(P)*), and rifamycin resistance (*rpoB2*), among others (*rsmA*, *arr-4*) (Figure S5b). Additionally, a noteworthy observation was the correlation of a considerable number of ARGs with multiple ROS stressors, specifically 24 ARGs correlating with *soxR*, *oxyR*, *sodAB*, and *katG* (including *AxyY*, *MuxC*, *OXA-145*, *mexY*, and others), and 11 ARGs correlating with *soxR*, *oxyR*, and *katG* (including *mtrA*, *vanD*, *mphA*, and others). Comparative analysis of both WWTPs revealed many common ARGs showing correlations with various ROS stressors, including genes encoding beta-lactamases (*OXA-4*, *PAU-1*), tetracycline resistance (*tet(T)*, *tetA(60)*), rifamycin resistance (*rpoB2*), and various multidrug resistance mechanisms (*adeF*, *AxyY*, *smeE*), suggesting shared patterns of ROS-mediated antibiotic resistance regulation across different wastewater treatment environments. These observed patterns align with and extend previous research findings in the field. For instance, studies with *Escherichia coli* K12 demonstrated that strains resistant to multiple antibiotics (chloramphenicol, amoxicillin and tetracycline) produced significantly higher levels of ROS compared to the wild-type (Jin et al., 2018). Additionally, the resistant strains also exhibited significant overexpression of ROS scavengers including *sodA* and *ahpF*. A recent study showed that exposure to bactericidal antibiotics (amoxicillin, enrofloxacin, and kanamycin) increased ROS production in the *oxyR* knocked out *E. coli* compared to the wild-type (Qi et al., 2023). Moreover, heme induced electron transport chain enhanced the ROS production, leading to *de novo* acquired resistance in *Lactobacillus lactis* (Qi et al., 2023). These reports support our findings on the pattern of correlations and underscore the potential link between ROS stressors and ARG emergence. A notable aspect of our findings is the identification of common and unique ARGs across different ROS stressors. Certain ARGs appear frequently across multiple ROS stressor correlations, suggesting that these ARGs might be more universally responsive to ROS-induced stress. In contrast, unique ARGs correlating with specific ROS stressors indicate targeted responses to particular oxidative stress conditions. Thus far, through meticulous examination of ROS stressor genes and their correlations with differentially enriched ARGs in the CAS aeration process, we uncovered patterns suggesting ROS as an instrumental factor in driving ARG enrichment/emergence.

### 3.4. Experimental support for increased ARG abundance and ROS under high DO

To investigate the mechanistic relationship between ROS and ARG enrichment/emergence within the CAS process, we conducted batch reactor experiments using CAS samples from the Māngere WWTP (Auckland, New Zealand). This experimental setup was designed to simulate the oxidative conditions characteristic of the CAS process, with specific emphasis on DO concentration variations. Our experimental design incorporated two distinct DO concentrations: DO2 (2 mg/L) and DO8 (8 mg/L), enabling assessment of ROS generation under elevated DO conditions. Quantitative analysis revealed mean H_2_O_2_ concentrations of 32.05 ± 2.08 μM and 38.38 ± 0.86 μM for DO2 and DO8 conditions, respectively (Figure 3a). The DO8 treatment yielded significantly higher H_2_O_2_ production compared to DO2 (p < 0.05). Analysis of oxidative stress response genes (*soxR*, *oxyR*, *sodAB*, *katG*, and *ahpCF*) demonstrated consistent upregulation under DO8 conditions (Figure 3b), indicating systematic adaptation to elevated ROS levels associated with higher DO concentrations. Resistome profiling revealed ARG patterns consistent with those observed in both WWTP_Gö_ and WWTP_Gw_ CAS systems (Figure S6a, b). The prevalent ARG classes encompassed resistance to aminocoumarin antibiotics, aminoglycosides, beta-lactams (including carbapenems, cephalosporins, penams), fluoroquinolones, macrolides, tetracyclines, peptide antibiotics, rifamycins, and sulfonamides (Figure S6c). Differential abundance analysis identified 11 significantly enriched ARGs under DO8 conditions (Figure 3c), including beta-lactamases (*OXA-34*), sulfonamide resistance (*sul4*), tetracycline resistance (*tet(43)*, *tet(W/32/O)*), rifamycin resistance (*Bado_rpoB_RIF*, *rphA*), and MDR (*mexQ*).

**Figure 3.**
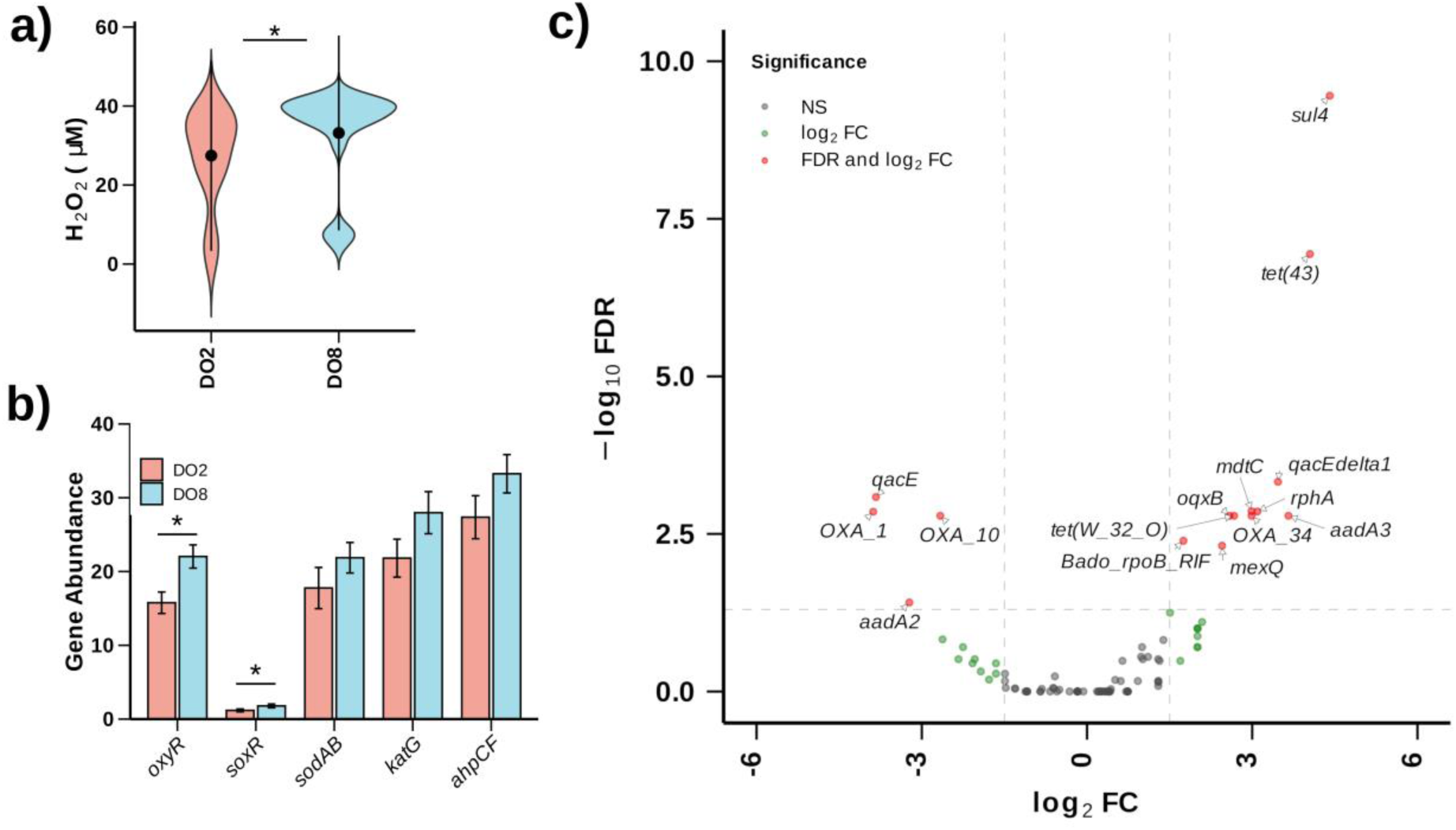
DO concentration influences ROS generation and ARG abundance. a) Violin plots showing H_2_O_2_ concentrations under DO2 (2 mg/L) and DO8 (8 mg/L) conditions (*p < 0.05, paired t-test). b) Relative abundance of oxidative stress response genes under DO2 and DO8 conditions. c) Volcano plot of differentially abundant ARGs comparing DO8 versus DO2 conditions (log_2_FC ≥ 1.5, FDR < 0.05).

To establish causality beyond correlation and verify protein expression, we performed untargeted SWATH metaproteomics analysis. Results demonstrated enhanced expression of several ARGs under DO8 conditions, including beta-lactamases (OXA-184, OXA-576, PME1), rifamycin resistance (RpoB2), tetracycline resistance (Tet(W/32/O)), and aminoglycoside resistance (AAC(6’)-Ia) (Figure 4). While some proteins showed elevated expression under DO2 conditions (Bpse_Omp38, K15582), others exhibited comparable expression levels across both conditions (BUT-1, JOHN-1, OXA-153). These findings provide mechanistic insights into ARG dynamics within CAS processes. The correlation between elevated ROS levels under high DO conditions and upregulation of oxidative stress response genes support that increased ROS production, resulting from intensive aeration in CAS systems, may contribute to ARG emergence and persistence.

**Figure 4.**
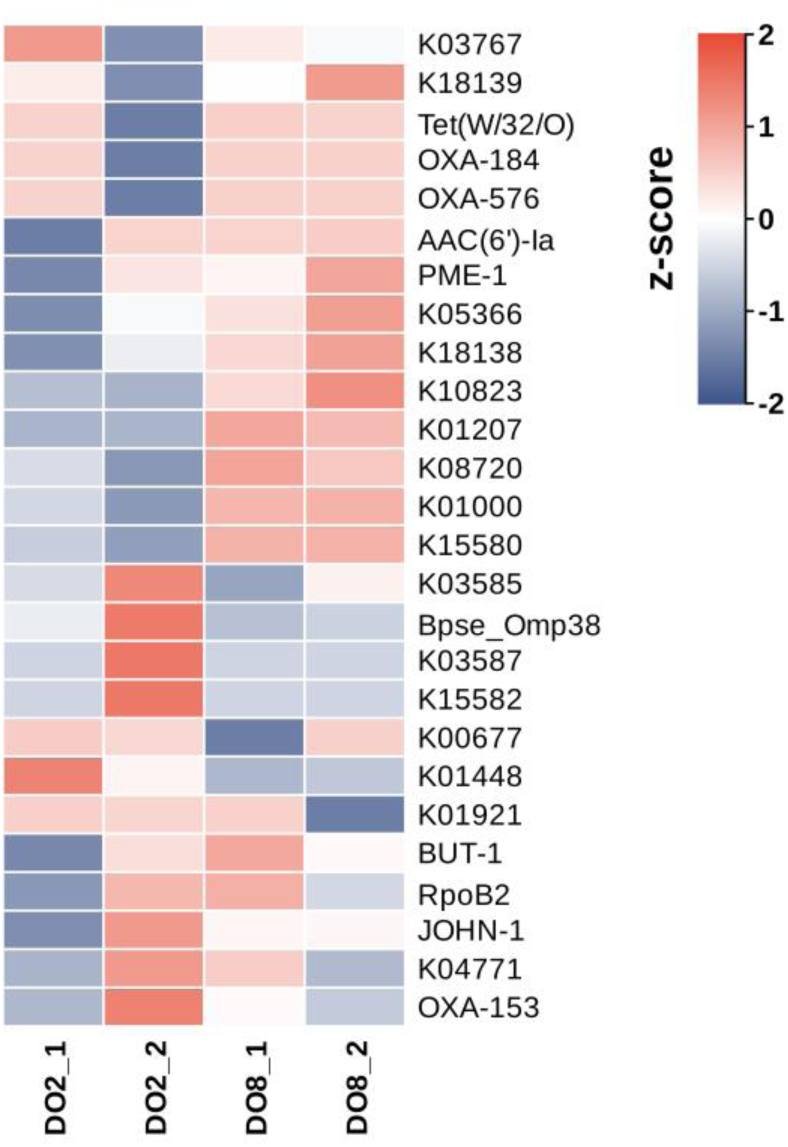
Heatmap with the protein expression of ARGs under DO2 and DO8 conditions measured by SWATH-MS based metaproteomics analysis. The protein names refer to the ARGs from CARD and KEGG orthologs.

### 3.5. Mechanism of ROS-induced ARG enrichment/emergence in CAS

Sublethal concentrations of endogenous ROS can trigger a series of direct or indirect cellular responses that may contribute to the enrichment/emergence of ARGs in the CAS process (Li et al., 2021). These responses include the activation of transcriptional regulators, expression of oxidative defense enzymes, metabolic adaptations, DNA damage, activation of efflux pumps, cell damage, and increased permeability and transformation potential (Figure S1). Firstly, enhanced expression of transcription factors like *oxyR* and *soxRS* facilitate the production of ROS-scavenging enzymes such as superoxide dismutase, catalase, thioredoxins and others (Imlay, 2008). In both the WWTP_Gö_ and WWTP_Gw_ CAS processes, we observed a high abundance of these transcription factors and ROS scavengers (Figure 5 a,b). Secondly, ROS stress can disrupt the TCA cycle, leading to hyperactivation of respiration, as seen in certain antibiotic treatments (Kohanski et al., 2008). Previous reports also suggested the disruption of the nucleotide pool, leading to the increase of ATP demand elevating TCA cycle activity and cellular respiration under quinolones treatments (Yang et al., 2019). This disruption may trigger metabolic adaptations, such as activating the glyoxylate shunt (GS) (Lemire et al., 2017) to reduce oxidative burden or redirecting flux to the PPP for increased NADPH production, supporting antioxidant defense systems (Christodoulou et al., 2018). Our comparative analysis revealed an higher abundance of genes involved in these metabolic pathways, ATP and NADPH production, and consumption in the CAS system compared to other treatment processes (Figure 5 a,b). Moreover, sublethal ROS levels can cause DNA damage and potential mutations, facilitating the emergence of *de novo* antibiotic resistance (Qi et al., 2023). Notably, DNA repair-related genes were abundant in the CAS processes at both WWTPs. We also identified several ARGs in the CAS process that were not detected in the upstream processes of both WWTPs (Supplemental Data 2b). This indicates that ROS-induced mutations could be responsible for their emergence. As a secondary defense, bacteria activate multidrug efflux pumps to expel antibiotics from the cell, promoting antibiotic tolerance. The abundance of multidrug efflux pump genes was high in the CAS processes (Figure 5 a,b). Additionally, oxidative damage can weaken the bacterial cell envelope, potentially leading to the release of ARGs into the environment. This process, coupled with enhanced transformation capabilities under oxidative stress, increases the uptake of foreign DNA, including ARGs, through HGT. Increased cell permeability can promote the uptake of foreign ARGs and eDNA through transformation (Y. Wang et al., 2020). Previous reports suggested that *Streptococcus* sp. released eDNA under H_2_O_2_ stress (Itzek et al., 2011). A recent report revealed that phenolic compounds can promote ROS-induced increased cell permeability and HGT (Ma et al., 2021). These simultaneous events occurring in the CAS process under oxidative stress conditions may play a significant role in enriching ARGs or helping ARG emergence. Overall, the proposed mechanisms by which ROS can contribute to ARG enrichment/emergence in the CAS process include activation of oxidative stress responses and metabolic adaptations, DNA damage and *de novo* resistance development, increased efflux pump activity, and enhanced cell permeability promoting horizontal gene transfer facilitated by oxidative damage. This multifactorial process underscores the complex interplay between oxidative stress and antibiotic resistance in the CAS process.

**Figure 5.**
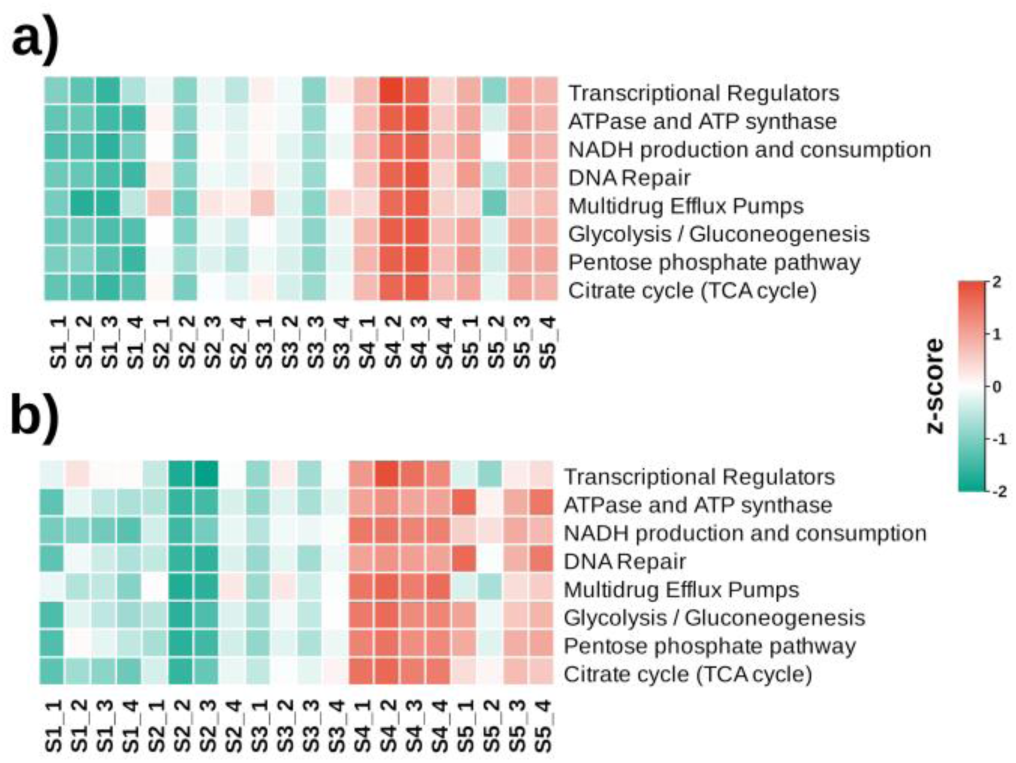
Heatmap illustrating the change in gene abundances associated with the cellular processes across various stages of the (a) WWTP_Gö_, and (b) WWTP_Gw_.

## 4. Conclusions

Our findings reveal a crucial paradox in CAS processes: while they effectively reduce overall ARG loads from hospital wastewater (retaining only 35-53% in effluent), they simultaneously create conditions that promote enrichment and emergence of specific ARGs. The demonstration that 30% of ARGs in CAS show higher abundance, with 22% of these enriched ARGs emerging *de novo*, challenges the traditional view of treatment efficiency. Beta-lactamases and efflux pumps, constituting nearly 50% of enriched ARGs, highlight the selective pressures within CAS systems. Most significantly, our finding that CAS resistome determines 58-75% of effluent ARG profiles indicates that treatment processes, rather than influent composition, shape the final resistome. The mechanistic link between ROS generation and ARG proliferation, validated through batch experiments, demonstrates how ancient oxidative stress responses continue to influence modern antibiotic resistance dissemination through engineered systems. Our findings highlight the need to consider AMR emergence as an integral component of the design and operation of biological wastewater treatment processes.

## Associated content

### Supporting Information

All the supplementary methods, tables and figures are provided in Supporting Information.pdf. Additionally, detailed information on differentially abundant ARGs, ARG family and drug classes, and ARG correlation with ROS genes are provided in Supporting Data.xlsx.

### Author information

CRediT authorship contribution statement

BM: Conceptualization, Investigation, Methodology, Formal Analysis, Writing - original draft. XZ: Investigation, Methodology, Writing - review & editing. NS: Conceptualization, Funding acquisition, Supervision, Writing - review & editing.

### Funding Sources

The study was funded by Marsden Award from the Royal Society of New Zealand (Contract Number: MFP-UOA2018) made to NS.

### Notes

The authors declare no competing financial interest.

## Acknowledgment

The authors acknowledge the use of New Zealand eScience Infrastructure (NeSI) high performance computing facilities, consulting support and/or training services as part of this research. New Zealand’s national facilities are provided by NeSI and funded jointly by NeSI’s collaborator institutions and through the Ministry of Business, Innovation & Employment’s Research Infrastructure programme. URL https://www.nesi.org.nz. The authors also acknowledge the Centre for eResearch at the University of Auckland for their help in facilitating this research. http://www.eresearch.auckland.ac.nz.

