## Supporting Information for "ROS-Induced Stress Promotes Enrichment and Emergence of Antibiotic Resistance in Conventional Activated Sludge Processes"

### for

Table of Contents Pages

Supplementary Methods, pages S2-S3

Tables S1-S5, pages S4-S8

Figures S1-S6, pages S9-S14

References, page 15

#### Supplementary Methods

##### ROS measurements

ROS level was assessed by measuring the hydrogen peroxide ($H_2O_2$) concentration with the Hydrogen Peroxide Sensor (ATT Bioquest, 11503) commercial kit. The fluorescence reaction system was set up in a 96-well plate, including 10 μL of 0.3% sodium azide, 50 μL of fluorescent probe working solution, and 50 μL of the sludge sample. Reactions were incubated at 20ºC, protected from light for 60 min, and fluorescence intensity was monitored with Ex/Em = 490/525 nm.

##### DNA extraction and metagenomics analysis

The procedure for DNA extraction started with collecting 10 mL sludge samples across DO2 and DO8 experimental conditions. The extraction followed the method provided by the DNeasy PowerSoil Kit (Qiagen, Germany), and the resultant DNA was preserved at -20°C for further analysis. The DNA samples were then sent off to the Auckland Genomics Centre (Auckland, NZ) to conduct metagenomic analysis. For all samples, a total amount of 500 ng DNA input was taken for library preparation. Libraries were made with the NEBNext® Ultra™ II DNA Library Prep Kit for Illumina® (E7645S) with unique dual indexes (cat no E7600). This process comprised several steps: Tagmentation of the samples to obtain a distribution of fragment lengths (200-450bp insert size) that are compatible with Illumina sequencing, ligation of Illumina adaptors to the fragment ends, performing a bead clean-up of the library to remove unligated adaptors and to impose some size selection (275-475bp insert size) on the library, incorporating the unique dual indexes by PCR, performing another bead clean-up to remove unincorporated indexes and excess primers, and QC libraries (including molarity estimation for pooling). Post library QC, normalization was carried out, followed by a final pool check using a bioanalyzer. The shotgun whole genome sequencing was carried out on a HiSeqX platform using 2x150 bp paired-end sequencing.

##### Untargeted metaproteomics analysis (SWATH)

Metaproteomic analysis was performed on sludge samples under various experimental conditions, using three biological replicates per condition. Sludge samples (5 mL) were pelleted via centrifugation at 6,113 x g for 5 min at 4°C and washed twice with 0.85% NaCl solution. The pellets washed were flash-frozen in liquid nitrogen and stored at -80°C. Protein extraction was conducted by resuspending the pellet in lysis buffer (pH 8) containing 2% (w/v) SDS, 2 mM EDTA, 2 mM phenylmethylsulfonyl fluoride, 0.15% Triton X-100, 20 mM DTT, and 50 mM HEPES and sonicating for 20 cycles with 15 seconds of on/off intervals on ice. The lysate was collected following centrifugation at 17,467 x g for 30 min at 4°C. The lysate was mixed with 20% trichloroacetic acid in acetone, vortexed, and incubated on ice. The mixture was centrifuged at 15,000 x g for 6 minutes at 4°C, and the supernatant was discarded. The pellet was washed with acetone, centrifuged, and washed again with 80% acetone in water. The protein pellet was air-dried and dissolved in 50 mM Tris buffer. Protein purification was further performed using SpeedBead carboxylate-modified E3 and E7 magnetic particles (Sera-Mag, USA). Protein concentration was quantified using the EZQ protein assay kit as per the manufacturer's instructions (Invitrogen, USA). A 250 µL sample containing 150 µg protein extract was reduced by adding 5 mM DTT and incubation at 56°C for 15 minutes. Alkylation was carried out by adding 15 mM iodoacetamide and incubating in the dark for 30 minutes. The reaction was quenched by adding 15 mM cysteine. Protein digestion was performed by adding 1.5 µg trypsin and incubating at 37°C for 1 hour, followed by 18 hours at room temperature. After digestion, the sample was diluted with 100 mM ammonium bicarbonate and centrifuged with a Vivaspin centrifugal concentrator (Sartorius, Germany) for 55 minutes at 28°C to intercept large molecules and obtain peptides filtered through. The sample was then diluted with 0.1% formic acid and peptides were extracted using Oasis Prime HLB 1cc (30 mg) solid-phase extraction. The peptide sample was loaded onto cartridges, which were subsequently washed with 1 mL of 0.1% formic acid. Peptides were eluted with 300 µL of 50% acetonitrile in 0.1% formic acid and concentrated to 15-20 µL using a speed vacuum. A 10 µL aliquot of each peptide sample was analyzed by nano LC-MS/MS. The sample was desalted and separated using a NanoLC 400 UPLC system (Eksigent, USA). The TripleTOF 6600 Quadrupole-Time-of-Flight mass spectrometer (Sciex, USA) was used for mass spectrometry analysis.

The resulting data were searched against a database established by metagenomics results of the samples using MetaProteomeAnalyzer version 3.4 (Muth et al., 2015). The X-tandem was selected as the primary peptides and protein identification search engine (Xu et al., 2013). The parameters were as follows: precursor and fragments tolerance set as 15 ppm; the cysteine alkylation set as iodoacetamide; the trypsin as the digestion enzyme with up to one mis-cleavage; a 1% false discovery rate was set as the filter of the final global protein groups. The resulting group file exported from X-tandem was converted to MetaProteomeAnalyzer for metaproteomics analysis and clustering.

**Table S1. Detailed sample statistics in WWTP$_{Gö}$.**

| WWTP Stages | Sample Name | NCBI SRA Accession | Number of Reads | Number of ORFs |
|---|---|---|---|---|
| **Hospital Effluent (S1)** | S1_1 | SRR11088470 | 50260226 | 3960819 |
| | S1_2 | SRR11088430 | 31474060 | 3366922 |
| | S1_3 | SRR11088412 | 64270110 | 3998257 |
| | S1_4 | SRR11088393 | 29595196 | 2864946 |
| **WWTP Influent (S2)** | S2_1 | SRR11088478 | 37916846 | 4058468 |
| | S2_2 | SRR11088438 | 31501626 | 3107742 |
| | S2_3 | SRR11088419 | 41526182 | 3541152 |
| | S2_4 | SRR11088401 | 38962344 | 3571546 |
| **Primary Treatment (S3)** | S3_1 | SRR11088471 | 35895430 | 3895742 |
| | S3_2 | SRR11088431 | 27028896 | 3631651 |
| | S3_3 | SRR11088413 | 19331728 | 2447689 |
| | S3_4 | SRR11088394 | 46792742 | 4319730 |
| **Activated Sludge (S4)** | S4_1 | SRR11088473 | 43040884 | 6014366 |
| | S4_2 | SRR11088434 | 90181452 | 8787764 |
| | S4_3 | SRR11088415 | 203935634 | 11628588 |
| | S4_4 | SRR11088396 | 33214154 | 6170529 |
| **WWTP Effluent (S5)** | S5_1 | SRR11088479 | 44235386 | 4780591 |
| | S5_2 | SRR11088439 | 32680206 | 4849985 |
| | S5_3 | SRR11088420 | 45461460 | 5531717 |
| | S5_4 | SRR11088402 | 33348936 | 5026031 |

**Table S2. Detailed sample statistics in WWTP Greifswald ($WWTP_{Gw}$).**

| WWTP Stages | Sample Name | NCBI SRA Accession | Number of Reads | Number of ORFs |
|---|---|---|---|---|
| **Hospital Effluent (S1)** | S1_1 | SRR11088387 | 49422328 | 3384143 |
| | S1_2 | SRR11088421 | 39038110 | 4006062 |
| | S1_3 | SRR11088403 | 48580442 | 4216584 |
| | S1_4 | SRR11088480 | 39029014 | 3792532 |
| **WWTP Influent (S2)** | S2_1 | SRR11088467 | 51321560 | 4443704 |
| | S2_2 | SRR11088427 | 13317556 | 2823495 |
| | S2_3 | SRR11088408 | 15743664 | 2696743 |
| | S2_4 | SRR11088390 | 76157828 | 6008780 |
| **Primary Treatment (S3)** | S3_1 | SRR11088388 | 19165630 | 3550005 |
| | S3_2 | SRR11088423 | 61889512 | 5360764 |
| | S3_3 | SRR11088404 | 30189644 | 3657943 |
| | S3_4 | SRR11088481 | 70784280 | 5486126 |
| **Activated Sludge (S4)** | S4_1 | SRR11088465 | 46516574 | 6036906 |
| | S4_2 | SRR11088425 | 55235376 | 6872171 |
| | S4_3 | SRR11088406 | 37323294 | 5580086 |
| | S4_4 | SRR11088483 | 51336318 | 6593468 |
| **WWTP Effluent (S5)** | S5_1 | SRR11088468 | 54264976 | 4008607 |
| | S5_2 | SRR11088428 | 57129998 | 4891412 |
| | S5_3 | SRR11088409 | 33412506 | 4023780 |
| | S5_4 | SRR11088391 | 41953798 | 4802367 |

**Table S3. Composition of artificial wastewater.**

| Chemicals | Concentration |
|---|---|
| Methanol | 26.936 mL/L |
| $NH_4Cl$ | 30.5712 g/L |
| $KH_2PO_4$ | 2.755 g/L |
| $K_2HPO_4$ | 2.755 g/L |
| $NaHCO_3$ | 76.8 g/L |

**Table S4. Composition of the trace element solution.**

| **Chemicals** | **Concentration (g/50 mL)** |
|---|---|
| EDTA | 0.125 |
| $ZnSO_4.7H_2O$ | 0.055 |
| $CoCl_2.6H_2O$ | 0.04 |
| $MnCl_2.4H_2O$ | 0.1275 |
| $MgSO_4.7H_2O$ | 1 |
| $CuSO_4.5H_2O$ | 0.043 |
| $(NH_4)_6Mo_7O_{24}.4H_2O$ | 0.0034 |
| $CaCl_2.2H_2O$ | 0.1375 |
| $FeSO_4.7H_2O$ | 0.1286 |

**Table S5. Detailed sample statistics in DO2 and DO8 experiments under the ENA Accession PRJEB74089.**

| WWTP Stages | Sample Name | ENA Accession | Number of Reads | Number of ORFs |
|---|---|---|---|---|
| **DO2** | DO2_1 | ERS18589221 | 10317308 | 1160874 |
| | DO2_2 | ERS18589222 | 12974262 | 1249751 |
| | DO2_3 | ERS18589223 | 17087718 | 1474762 |
| | DO2_4 | ERS18589224 | 19425152 | 1498240 |
| | DO2_5 | ERS18589225 | 12509546 | 1178711 |
| **DO8** | DO8_1 | ERS18589238 | 30287196 | 1813726 |
| | DO8_2 | ERS18589239 | 29660950 | 1795808 |
| | DO8_3 | ERS18589240 | 19906632 | 1468815 |
| | DO8_4 | ERS18589241 | 28489244 | 1699436 |

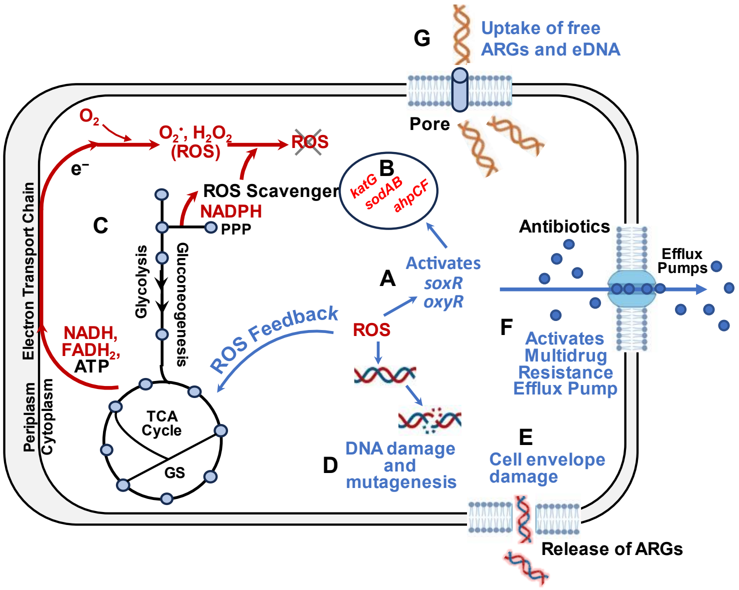


**Figure S1.** Mechanisms of ROS-induced ARG enrichment/emergence. a) A schematic illustration of the possible mechanisms by which ROS may contribute to ARG proliferation in the CAS process. The diagram depicts the cascade of cellular responses triggered by ROS, including (A) activation of transcription factors that (B) induced expression of ROS-scavenging enzymes, (C) metabolic adaptations to mitigate ROS damage, (D) DNA damage leading to mutations, (E) cell envelope damage and ARG release, (F) activation of multidrug resistance efflux pumps, (G) uptake of free ARGs and eDNA. These responses collectively facilitate an environment conducive to the proliferation and spread of ARGs in the CAS system. Adopted from Li, Hao, et al (Li et al., 2021).

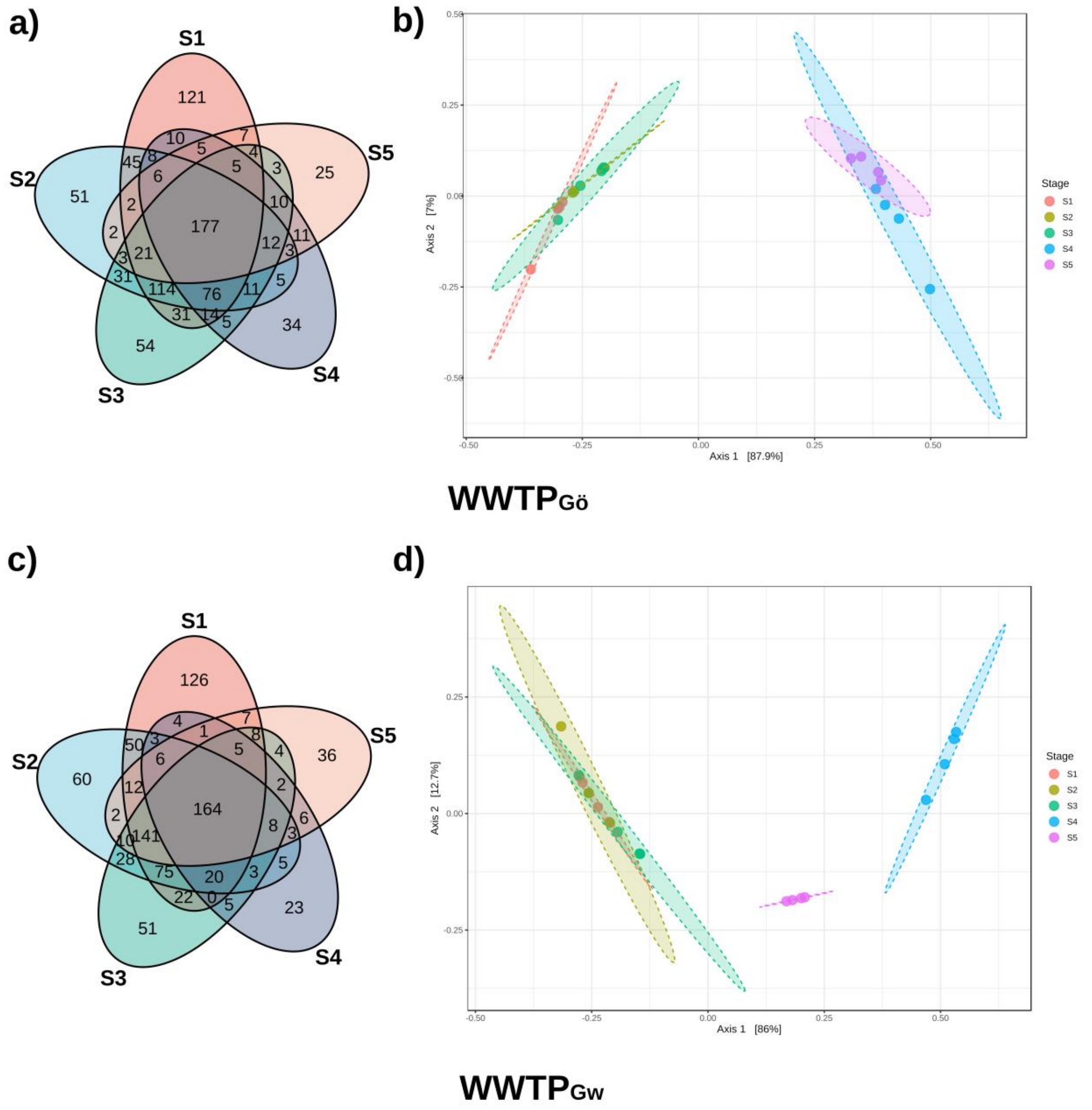


**Figure S2.** Venn diagram showing the shared ARGs across different treatment stages in a) $WWTP_{Gö}$., and c) $WWTP_{Gw}$. Principal Coordinates Analysis (PCoA) plot illustrating the beta-diversity of the ARGs in b) $WWTP_{Gö}$., and d) $WWTP_{Gw}$.

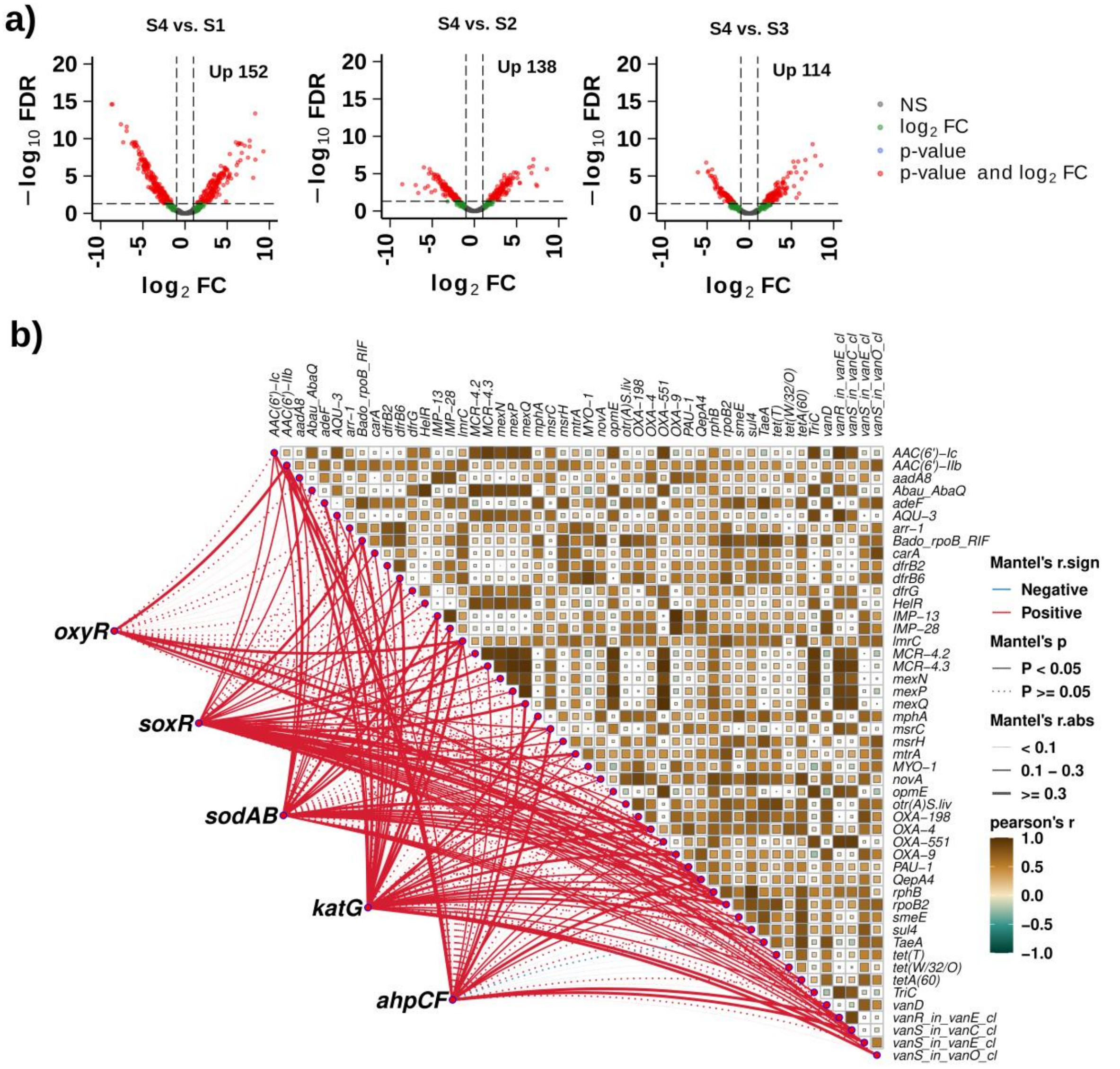


**Figure S3.** Differentially enriched (or emerged) ARGs and their correlation network with ROS stressors in the CAS process. a) Volcano plots illustrating the differentially abundant ARGs in the CAS stage compared to S1: Hospital effluent, S2: WWTP influent, S3: Primary sludge in the $WWTP_{Gö}$ ($\log_2 FC \geq 1$, FDR < 0.05). b) A network analysis depicting the correlations between selected enriched/emerged ARGs and ROS stress response genes in CAS process in $WWTP_{Gö}$.

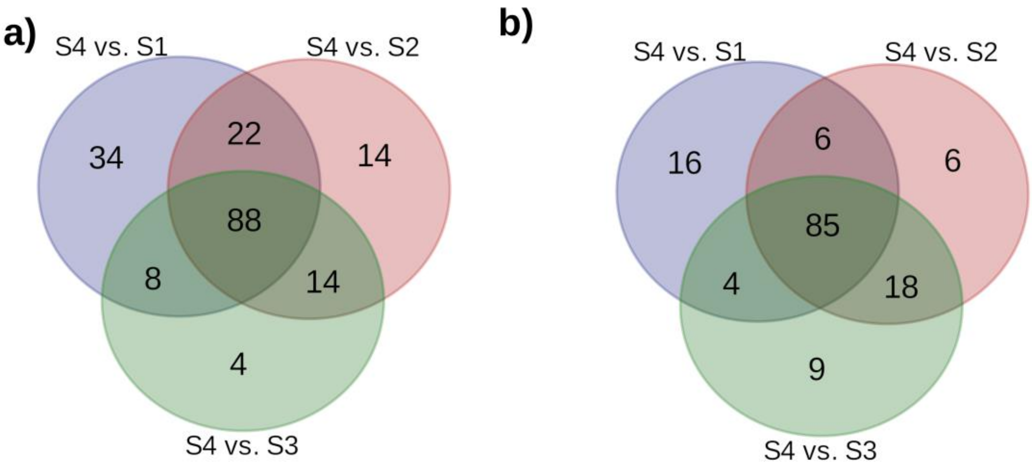


**Figure S4.** Venn diagram of the number of differentially higher abundant ARGs in CAS (S4) compared to hospital effluent (S1), WWTP influent (S2), and primary sludge (S3) in a) $WWTP_{Gö}$, and b) $WWTP_{Gw}$.

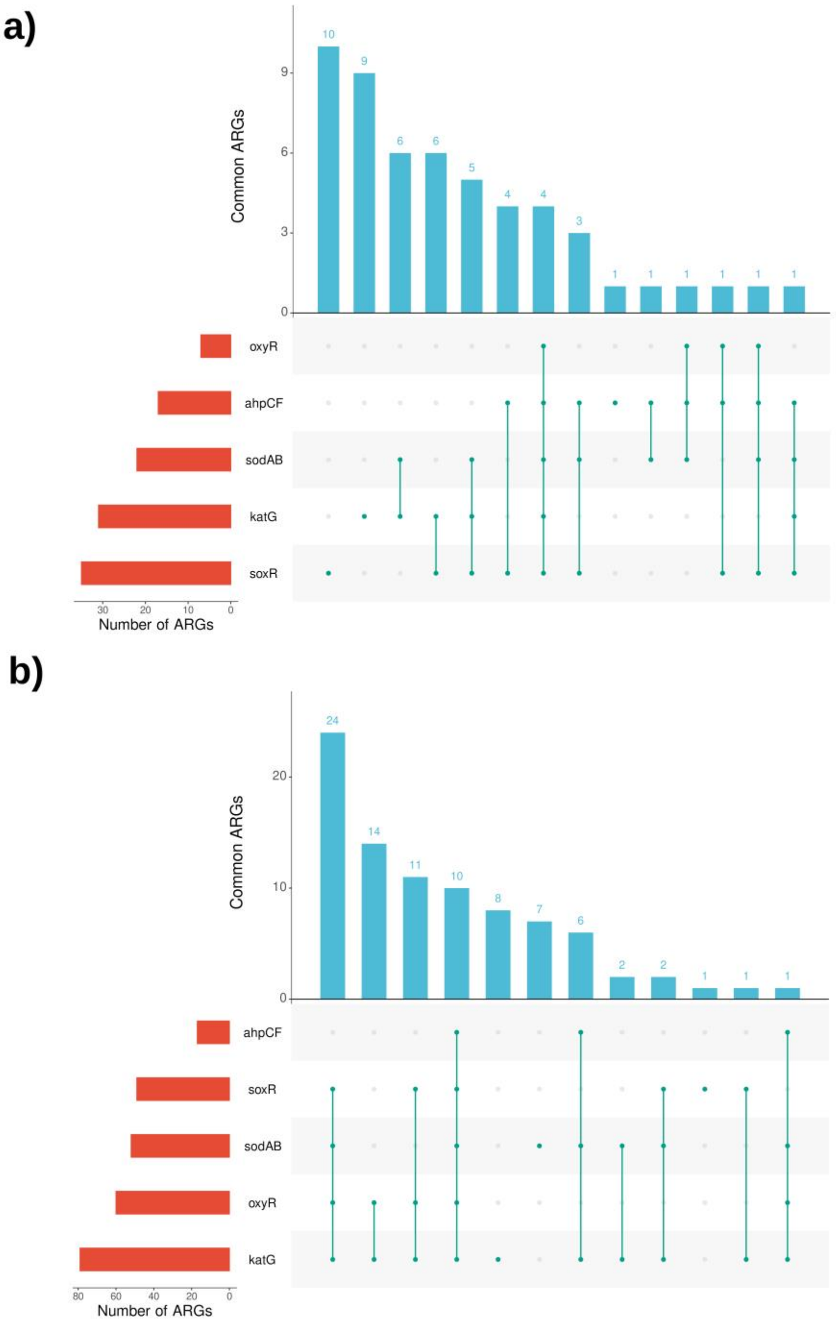


**Figure S5.** Upset diagram showing the number of enriched/emerged ARGs positively correlated with the ROS stressors in a) WWTP$_{Gö}$, and b) WWTP$_{Gw}$.

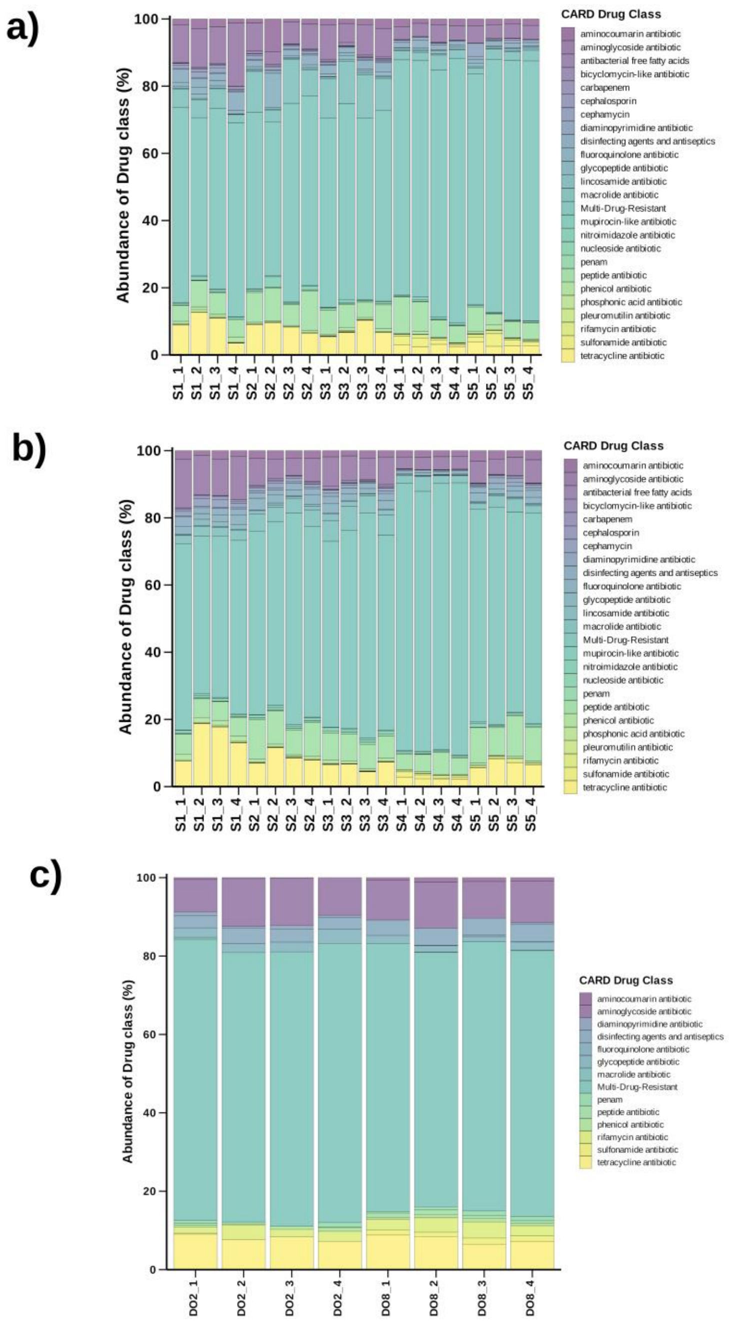


**Figure S6.** Stacked bar plot showing the relative abundance of ARGs resistant to different antibiotic drug classes in a) WWTP$_{Gö}$, b) WWTP$_{Gw}$, and c) DO (DO2 vs. DO8) experiments.
